# Data driven shoe design improves running economy beyond state-of-the-art Advanced Footwear Technology running shoes

**DOI:** 10.1101/2025.04.13.648601

**Authors:** John Kuzmeski, Montgomery Bertschy, Laura Healey, Zach Barrons, Wouter Hoogkamer

**Author notes:** **Corresponding author** John Kuzmeski. **Author contributions** Conceptualization: John Kuzmeski, Montgomery Bertschy, Laura Healey, Zach Barrons, Wouter Hoogkamer; Experimental design: John Kuzmeski, Montgomery Bertschy, Zach Barrons, Wouter Hoogkamer; Data collection: John Kuzmeski, Montgomery Bertschy, Zach Barrons; Data analysis: John Kuzmeski, Montgomery Bertschy, Zach Barrons; Writing - original draft preparation: John Kuzmeski, Montgomery Bertschy, Laura Healey, Wouter Hoogkamer; Writing - review and editing: John Kuzmeski, Montgomery Bertschy, Laura Healey, Zach Barrons, Wouter Hoogkamer. **Funding** PUMA SE provided the shoes and funding for this project. **Disclosure statement** Wouter Hoogkamer has received research grants from PUMA and Saucony. Laura Healey is an employee of PUMA. PUMA did not have any influence on the views presented in this article.

## Abstract

Advanced Footwear Technology (AFT) has enabled remarkable improvements in running performance over traditional marathon racing shoes. However, reported differences between state-of-the-art AFT models are small, and vary across individuals. To assess if the benefits of AFT have been fully realized or if further running economy improvements can be unlocked using modern computational design and optimization techniques, we compared a prototype AFT shoe developed using a data-driven computational design process (PUMA Fast-R 3; FR3) against state-of-the-art AFT models. We quantified running economy for 15 trained runners (11M, 4F) in this prototype AFT shoe and three commercially available AFT models: the PUMA Fast-R 2 (FR2), the Nike Alphafly 3 (NIKE), and the Adidas Adios Pro Evo (ADI). Running economy in the FR3 was 3.15 ± 1.24%, 3.62 ± 1.25%, and 3.54 ± 1.16% better than in the FR2, NIKE and ADI (all p < 0.001), respectively, and every individual performed best in the FR3 shoes. While step parameters were similar between FR3 and FR2, the FR3 had a lower step frequency than the ADI (p = 0.013) and longer contact time than the NIKE and ADI (both p<0.001). Our results suggest that computational design analysis is a promising frontier in performance running shoe design, offering potential for further improvements and personalized AFT models.

## Introduction

In the past decade, running shoe innovation has led to exciting improvements in running performance^1–5^. Hoogkamer et al.^6^ were the first to study a novel type of marathon racing shoe featuring a modern, compliant and resilient foam and an embedded carbon fiber plate. They observed running economy improvements of 4% over traditional marathon racing shoes. Shoes with these and similar features are commonly referred to as “super shoes”, and as Advanced Footwear Technology (AFT) among researchers^7^. Since then, almost all running shoe brands have developed their own AFT model targeting improved running economy and performance. The individual contributions of reduced mass, increased longitudinal bending stiffness from carbon fiber plates, increased compliance and resilience from modern foams, taller stack height, and pronounced rocker profile are still topic of debate^8–14^, but virtually all state-of-the-art marathon racing shoes leverage some combination of these properties.

While several studies have evaluated and compared the first AFT model (Nike Vaporfly 4%) in different groups and under different conditions^15–23^, less have evaluated its successor (Nike Vaporfly next%)^24–28^, or other more recent state-of-the-art marathon racing shoes^29–34^. Insights from this body of literature suggest that the best AFT models have on average 3-4% better running economy than traditional marathon racing shoes, but that running economy differences between state-of-the-art AFT models are 1-1.5% on average, at best. For example, Denis et al.^29^ compared running economy between Adidas Adios Pro 3 and 4 models, released in June 2022 and January 2025 respectively, and observed a significant running economy improvement of 1.3% on average. This plateau in running economy gains over recent years raises the question of whether the benefits of AFT have been fully realized or if further improvements can be unlocked using modern computational design and optimization techniques.

Advancements in computational modeling and simulation provide product developers with powerful tools to virtually design, evaluate, and optimize a product before physical prototyping. Not only can such computational tools significantly reduce development time and resource costs, but they also enable a more comprehensive understanding of how the various design features influence product performance. In the case of footwear design, this process would typically begin with the creation of a parametric computer-aided design (CAD) model of a shoe or one of its components^35^. The CAD model is then converted into a parametric finite element model (FEM). Finite element analysis (FEA) is used to simulate forces experienced during running and evaluate key performance metrics^35^. Optimization algorithms can then iteratively tune design parameters of the FEM to meet specific performance goals (e.g., minimize mass and maximize energy return). While such techniques are commonly used in various engineering domains, it remains unclear to what extent these methods are being applied in the practice of footwear design, as companies seldom disclose their internal design workflows.

Here, we compared running economy between three state-of-the-art AFT models and a prototype shoe model developed using data-driven computational design analysis. We hypothesized that running economy would be better in the prototype than in the commercially available state-of-the-art AFT models and anticipated no detectable differences between the existing AFT models.

## Methods

### Shoe conditions

The prototype shoe (PUMA Fast-R 3) was developed by PUMA. To start, a virtual shoe model was created using the known geometry and material specifications of the PUMA Fast-R 2. Biomechanical data were recorded for 10 runners running at 14-18 km/h in the PUMA Fast-R 2. This data included time series of in-shoe pressure data (XSENSOR Technology Corporation, Calgary, Canada), ground reaction forces (Treadmetrix, Park City, UT, USA), and 3D kinematics from the ankle and metatarsal-phalangeal joint (Cameras: OptiTrack Color Prime, NaturalPoint Inc., Corvallis, OR, USA; Software: Theia 3D, Theia Markerless Inc., Kingston, Canada). These biomechanics data were used to simulate how the shoe behaves during running. Using dynamic computational simulations with the runners’ data, PUMA applied topology optimization to identify the optimal foam placement, material properties, carbon fiber layering, and overall geometry to minimize weight, maximize energy storage and return, and achieve the desired carbon plate stiffness.

Upon receiving the final prototype shoe from PUMA, we compared it against three state-of-the-art, commercially available AFT models, without further involvement of PUMA. We included the Nike Alphafly 3 (NIKE), Adidas Adios Pro Evo 1 (ADI), and PUMA Fast-R 2 (FR2) (Figure 1). At the start of the testing phase the NIKE and ADI shoes were the shoes worn for the male and female marathon world records, respectively.

**Figure 1.**
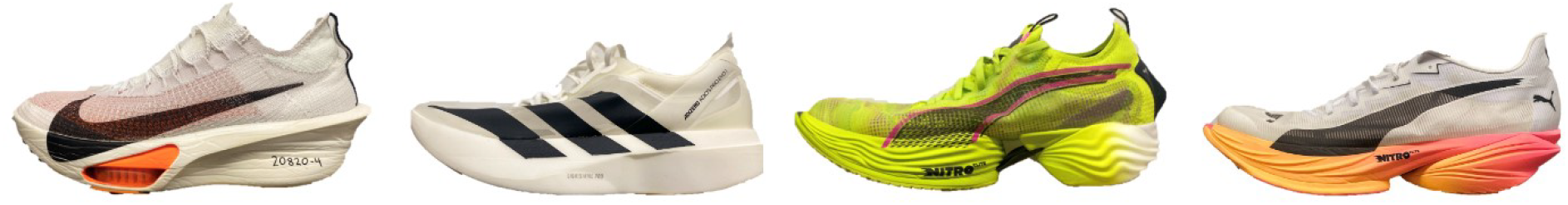
From left to right: Nike Alphafly 3 (NIKE), Adidas Adios Pro Evo 1 (ADI), PUMA Fast-R2 (FR2), PUMA Fast-R 3 (FR3).

### Material properties

We tested each shoe model’s material properties with an Instron E10000 (Instron, Canton, MA, USA). We used a custom WaveMetrix script to control two different missile heads, a flat round missile with a diameter of 45mm to mimic the size of the calcaneus for rearfoot testing and a full foot last with heel to toe drop of 10 mm to assure even contact across the midsole. Each missile head was force-controlled to mimic the peak force for the corresponding region of the shoe: 1200 N for the rear foot and 2000 N for the full foot. Additionally, each time period was adjusted to mimic the loading duration for each area based on insole data from Honert et al.^36^: 140 ms for the rearfoot and 185 ms for the full foot. Each shoe was subjected to 100 half sine wave load cycles, with the final 10 load cycles averaged for analysis.

Longitudinal bending stiffness was tested through a 3-point bending test. Each shoe was placed upon two parallel supports located^37^. The Instron tip was aligned with the metatarsal phalangeal joint, in the center of the supports, and followed a position controlled trapezoidal waveform while force was recorded^37^. This waveform depressed 7.5 mm into the midsole at 10 mm/s, was held that position for 0.5 seconds and then raised 7.5 mm at 10mm/s. This cycle was repeated 20 times, with the final 10 load cycles averaged for analysis. Shoe properties and mechanical testing results are summarized in Table 1.

**Table 1.**
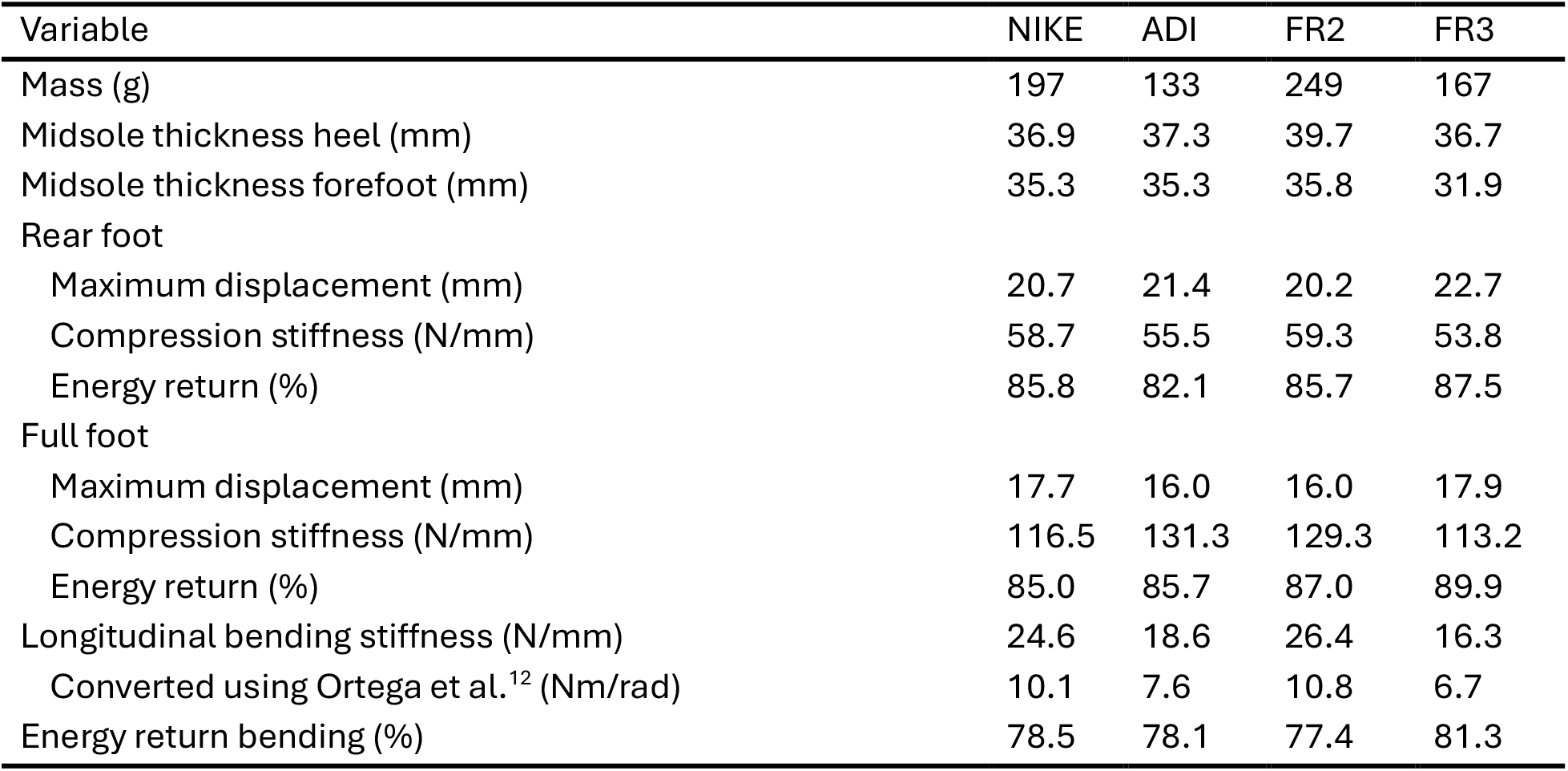
Material properties and mechanical testing results for the Nike Alphafly 3 (NIKE), Adidas Adios Pro Evo 1 (ADI), PUMA Fast-R 2 (FR2), and PUMA Fast-R 3 (FR3) for the men’s US 9 sized shoes.

**Table 2.** Spatiotemporal and kinetic outcomes between the four shoe conditions. NIKE: Nike Alphafly 3, ADI: Adidas Adios Pro Evo 1, FR2: PUMA Fast-R 2, FR3: PUMA Fast-R 3. * denotes significant effect of footwear condition.

| Variable | NIKE | ADI | FR2 | FR3 |
| --- | --- | --- | --- | --- |
| Step frequency (steps/min) * | 179.6 ± 9.1 | 182.0 ± 9.4 | 178.7 ± 8.4 | 179.6 ± 8.9 |
| Contact time (s) * | 0.210 ± 0.013 | 0.207 ± 0.012 | 0.215 ± 0.013 | 0.214 ± 0.012 |
| Peak vGRF (BW) | 2.68 ± 0.38 | 2.65 ± 0.41 | 2.70 ± 0.37 | 2.68 ± 0.38 |
| Braking & propelling impulses (BW•s) * | 0.024 ± 0.003 | 0.024 ± 0.003 | 0.025 ± 0.003 | 0.025 ± 0.003 |

### Participants

Fifteen runners participated in this study, 11 males (age: 31.5 ± 6.8 years old, height: 1.78 ± 0.04 m, mass: 67.5 ± 4.8 kg) and 4 females (age: 28.3 ± 3.4 years old, height: 1.66 ± 0.01 m, mass: 59.1 ± 2.2 kg). To be included in the study, participants were required to have recently run a 5-km in less than 21 minutes (females) or 19 minutes (males) or an equivalent race result over a different distance, and to fit a shoe size US W8, M9 or M11. The study was approved by the University of Massachusetts Amherst Institutional Review Board (#5924). All participants signed written informed consent forms before participating.

### Running Protocol

Upon arriving at the lab for testing participants warmed up as they would for a high tempo workout, on a treadmill or outside. Once fully warmed up participants ran on a rigid force instrumented treadmill (Treadmetrix, Park City, UT, USA) for 2 minutes to familiarize themselves with the treadmill and mouthpiece. For the experimental protocol, the participants ran in each of the four shoe models twice in randomized, mirrored order (ABCDDCBA) for 5 minutes, followed by 5 minutes rest, for a total of eight running trials, while we measured expired air (TrueOne 2400, ParvoMedics, Salt Lake City, UT, USA). Running speed for each runner was set based on experience and recent 5k race results. Eight males ran at 16 km/h, three males ran at 14 km/h, three females ran at 14 km/h, one female ran at 12.9 km/h (7:30 min mile pace). The expired gas analysis system was calibrated before the start of the first trial. We recorded ground reaction force data at 1000 Hz during the last 30 s of each trial.

### Data analysis

We calculated running economy as average metabolic rate (W/kg) over the last 2 minutes of each trial, based on the measured rates of oxygen uptake and carbon-dioxide production^38^. We filtered ground reaction force data using a dual-pass Butterworth filter with a 20 Hz cut-off^16,39^. Touch down and toe off were determined using a 25 N vertical ground reaction force threshold to calculate ground contact time and step frequency. We calculated peak vertical ground reaction force, and braking and propelling impulses during the contact phase.

### Statistics

We used linear mixed-effects models to evaluate the effects of shoe model on running economy and kinetic outcomes. We used a traditional level of significance (α = 0.05), with Tukey’s HSD test corrections on post hoc analysis. Shoe conditions were used as the fixed effect, participants as a random intercept, and running economy and kinetic outcomes each as output variable.

## Results

We observed a significant effect of shoe type on running economy (Fig. 2). Post-hoc pairwise comparisons, revealed that the FR3 had a significantly better running economy (15.82 ± 1.72 W/kg) compared to the FR2 (16.33 ± 1.67 W/kg; p < 0.001), NIKE (16.42 ± 1.83 W/kg; p < 0.001), and ADI (16.41 ± 1.86 W/kg; p < 0.001). For all 15 participants running economy was best in the FR3. The average percent running improvements in the FR3 were 3.15 ± 1.24%, 3.62 ± 1.25%, and 3.54 ± 1.16% vs. the FR2, NIKE and ADI, respectively. There were no significant differences in running economy between the FR2 and ADI (p = 0.515), FR2 and NIKE (p = 0.406), or NIKE and ADI (p = 0.998). The respiratory exchange ratio was below 1.0 for all participants during all trials (0.92 ± 0.02).

**Figure 2.**
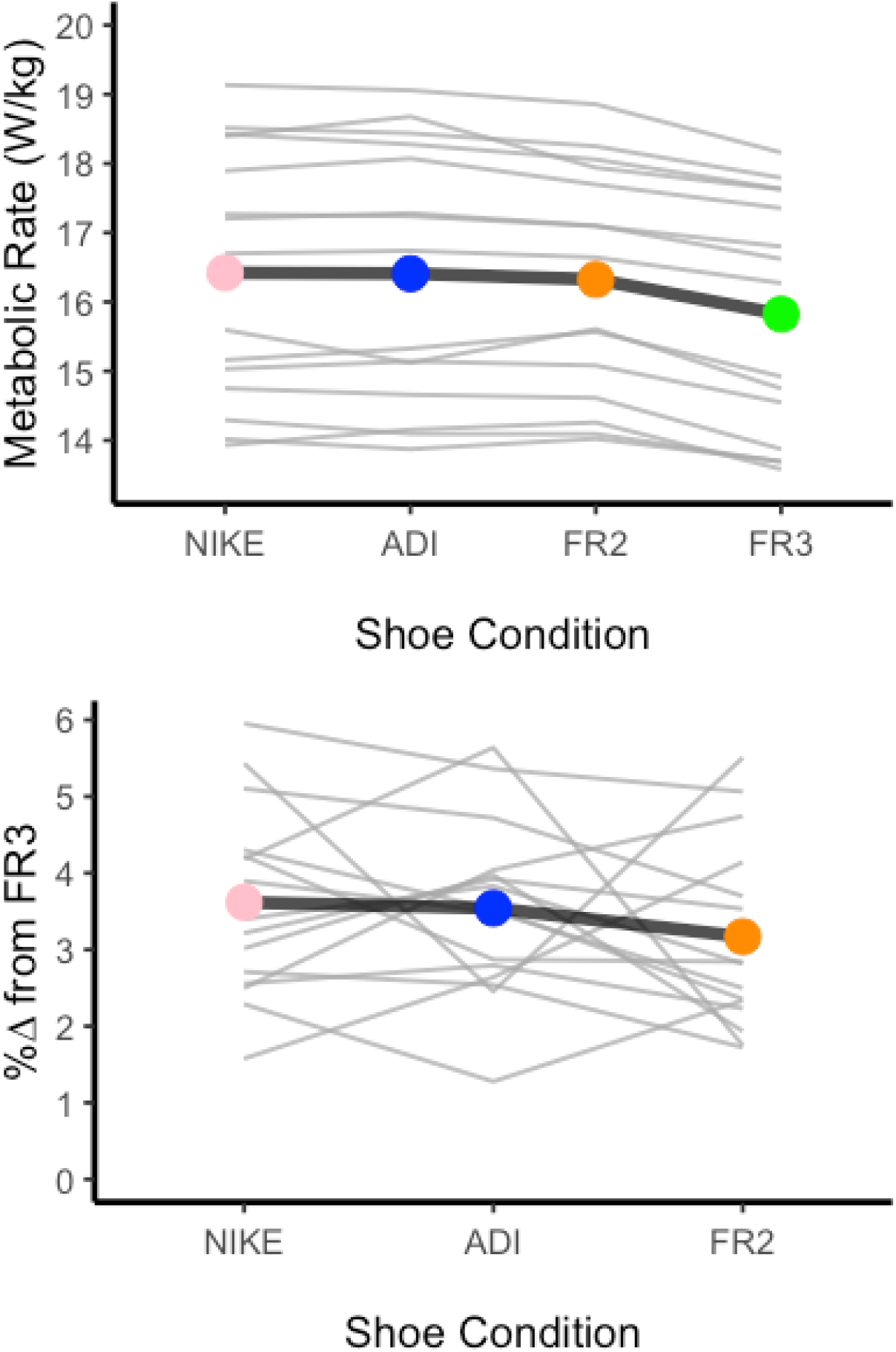
A) Running economy, quantified in metabolic rate, was best in the PUMA Fast-R 3 (FR3) vs. the Nike Alphafly 3 (NIKE), Adidas Adios Pro Evo 1 (ADI), and PUMA Fast-R 2 (FR2). B) Percent improvement in the FR3 relative to the other shoes varied substantially between individuals and shoes.

There was a significant effect of shoe on step frequency (p < 0.001), contact time (p < 0.001), and braking/propelling impulses (p < 0.001). Step frequency was higher in the ADI than in the FR2 (p < 0.001) and FR3 (p = 0.013), but not than in the NIKE (p = 0.083). It was similar for the other between shoe comparisons. Contact time was similar in the FR2 and FR3 (p = 0.763), both longer than in the ADI and NIKE (all p < 0.001). ADI and NIKE were not significantly different from each other (p = 0.154). Braking and propelling impulses were higher in the FR3 than in the ADI (p < 0.001) and NIKE (p = 0.024), but not than in the FR2 (p = 0.794). They were also higher in the FR2 than in the ADI (p = 0.002), but not than in the NIKE (p = 0.170). They were similar in ADI and NIKE (p = 0.327).

## Discussion

The primary finding of this study supports our hypothesis: the prototype shoe (FR3), developed through a data-driven computational design process, yields significantly better running economy compared to three commercially available state-of-the-art AFT models (NIKE, ADI, FR2). Specifically, running in the FR3 resulted in an average group improvement in running economy between 3.1 and 3.6% in relation to these state-of-the-art models, with each participant having the best running economy in the FR3. Furthermore, there was no significant difference in running economy between each of the commercially available state-of-the-art models with variation across participants in which of these shoes yielded the second-best running economy.

The most striking result from this study was the consistency of the running economy response to the prototype shoe (Fig. 2). Every participant recorded their best running economy while running in the FR3. While a similar consistent improvement has been reported for AFT before, this was relative to traditional racing shoes^6,15^. Others have reported inconsistent responses for running economy in AFT vs. traditional racing shoes, with some individuals showing no improvements or slightly worse running economy in AFT^19,20^. Running economy differences between AFT models are typically small and inconsistent between participants^29,31,32^. A common explanation for these inconsistencies is that there are “responders” and “non-responders”^40–42^. The unanimous positive response to the FR3 in our population highlights a further advancement in running economy for racing shoes, a step similar to the introduction of AFT, suggesting computational design analysis is a promising new frontier in performance running shoes design.

The observed between shoe differences in running economy were not mirrored by changes in spatiotemporal and kinetic variables. Participants ran with a lower step frequency in the FR3 than in the ADI, while contact time and braking/propelling impulses were larger in the FR3 than in the ADI and NIKE. Longer contact times could partially explain the improved running economy^43^. However, FR3 also led to a significantly improved running economy over its predecessor (FR2) despite no spatiotemporal or kinetic differences.

While the difference between traditional marathon racing shoes and AFT in appearance and properties is striking, the differences between the FR3 and the three other shoes is much more subtle. The computational design process focused on optimizing footwear characteristics known or expected to improve running economy such as mass, carbon fiber plate geometry and stiffness, and foam properties and distribution for maximal mechanical energy storage and return. While others have combined shoe finite element analysis with musculoskeletal models to estimate bone loading^44^ or have used musculoskeletal simulations to predict differences in running economy from different midsole foams^45^, the current process only focused on shoe performance. Through the optimization process, the mass of the FR3 (167g) came out 82 g lighter than the FR2, which is relatively heavy (249g) compared to the competitor AFT models (NIKE 197g, ADI 133g). The 1% per 100g rule^46,47^ indicates a ∼0.8% better running economy in the FR3 than in the FR2, which leaves ∼2.3% improvement in running economy to be accounted for by improvements in plate geometry and stiffness and foam energy storage and return.

Moreover, the FR3 is substantially less stiff in longitudinal bending than the FR2 (16.3 vs. 26.4 N/mm). Since we assessed this with a three-point bending test, part of the compliance comes from foam deformation rather than plate bending^48^. Compression stiffness was lower in the FR3 than in the FR2 for both rear foot and full foot compression. Percent energy return in the FR3 was higher than in the FR2, both in compression and in bending. While not the lightest, as compared to the three other shoes, the FR3 had the lowest compression stiffness, the lowest bending stiffness, and the highest percent energy return, suggesting that the observed improved running economy is a combination of all of these factors.

While all participants had their best running economy in the FR3, running economy in the three state-of-the-art AFT models varied substantially between individuals. The largest difference among the three AFT models was 4.1% for one individual (ADI vs. FR2), and across individuals running economy relative to the FR3 varied from 1.6-6.0% in NIKE, 1.3-5.6% in ADI and 1.7-5.5% in FR2. Such inter-individual variability is in-line with other studies using multiple trials in each shoe to assess running economy difference, report running economy improvements beyond traditional racing shoes ranging from ∼1% to ∼7%^6^ (for review see ^49^), but smaller than controversial findings showing >10% improvements and deteriorations in running economy between AFT models for some individuals ^49,50^. This 4 - 5% range in individual running economy values in the same shoes and the observed individual changes between the AFT models, confirms that some runners might have more benefits from specific AFT models than other runners, supporting the potential of personalized AFT design^51,52^. Here we show that AFT shoes developed using a data-driven computational design process improve running economy beyond these interindividual differences, as all participants had their best running economy in the FR3, but we see great potential for using data-driven computational design processes to optimize AFT for an individual runner.

Running economy is one of the three major determinants of marathon performance^53,54^, with recent insights suggesting a role for physiological resilience as well ^55^. Indeed, improvements in running economy translate to improvements in running performance, but this relation is not one-to-one^56^. Kipp et al^57^ quantified this relation accounting for the inherent curvilinear relation between running speed and running economy^58,59^, and the additional metabolic cost of overcoming air resistance during overground running, which increases disproportionally with increasing speed^60^. This relation indicates that the actual performance benefits from an improvement in running economy (e.g., time saved in a race) depend on the running velocity.

Specifically, at faster running speeds the percent time savings are less than the percent improvement in running economy, which is also what is commonly reported when analyzing race result improvements since the introduction of AFT^1–5^. To put the observed running economy improvements in context, we used the framework by Kipp et al.^57^ to calculate the theoretical improvements in marathon time from a 3.15% improvement in running economy (FR3 vs. FR2) for different baseline marathon times, for a sample runner (mass 65.6 kg and height 1.75 m)^61,62^ (Table 3). Future race result analyses are needed to evaluate these predicted time improvements.

**Table 3.** Predicted marathon time improvements from a 3.15% improvement in running economy for different baseline marathon times.

| Marathon time<br>[hr:min:sec] | Running economy<br>improvement [%] | Running time<br>improvement [%] | New marathon time<br>[hr:min:sec] |
| --- | --- | --- | --- |
| 2:00:00 | 3.15 | 2.04 | 1:57:36 |
| 2:30:00 | 3.15 | 2.30 | 2:26:37 |
| 3:00:00 | 3.15 | 2.61 | 2:55:26 |
| 3:30:00 | 3.15 | 2.94 | 3:24:00 |
| 4:00:00 | 3.15 | 3.32 | 3:52:17 |
| 4:30:00 | 3.15 | 3.73 | 4:20:18 |
| 5:00:00 | 3.15 | 4.17 | 4:48:00 |

Note that we only evaluated running economy for runners running at 12.9 (n = 1), 14 (n = 6) and 16 (n = 8) km/h, which limits extrapolation of our findings to elite runners (18km/h or faster) and recreational runners (12 km/h or slower)^27^. Furthermore, these footwear conditions may have a different effect if the participants are able to adapt over the course of several days or weeks, however all participants were used to regularly running in AFT models. By design, we compared state-of-the-art AFT models as is, with many differences in foam and plate properties, geometry and mass. As such our findings provide limited insights into the exact features that explain the observed differences in running economy. It might be that using more trials per condition (e.g., four) would have resulted in significant running economy differences between the three commercially available AFT models^63^, or that a larger sample size would have resulted in additional significant between shoe differences in spatiotemporal measures (e.g., step frequency in NIKE vs. ADI). Finally, we only assessed running economy over 5-minute trials and while changes in lab-based measures of running economy translate to running performance as discussed above, we did not assess how these shoes might compare under fatigued conditions to be anticipated 35 kilometers into a marathon. Similarly, we assessed shoes that were relatively fresh, used less than 40 km use per pair, we did not assess how these shoes compare after substantial use^64^.

## Conclusion

To assess if the benefits of AFT have been fully realized, we compared a prototype AFT shoe developed using a data-driven computational design process (FR3) to commercially available AFT models. The FR3 was developed starting from a virtual shoe model with the known geometry and material specifications of the FR2. Time series of in-shoe pressure data, ground reaction forces, and ankle and metatarsal-phalangeal joint kinematics from 10 runners were used to simulate how the shoe behaves during running and topology optimization was applied to identify the optimal foam placement, material properties, carbon fiber layering, and overall geometry to minimize weight, maximize energy storage and return, and achieve the desired carbon plate stiffness. Running economy improved over 3% in the FR3 vs. the FR2 and two competitor AFT models (NIKE and ADI). While step parameters were similar between FR3 and FR2, the FR3 had a lower step frequency than the ADI (p = 0.013) and longer contact time than the NIKE and ADI (both p<0.001). Our results suggest that computational design analysis is a promising frontier in performance running shoe design, offering potential for further improvements and personalized AFT models.

## Acknowledgements

We thank the subjects for participating, Joshua Cohen and Caroline Kolmodin for helping with data collection, and Dr. Meghan Huber and Dr. Key Nahan for proofreading this manuscript.

## Author contributions

Conceptualization: John Kuzmeski, Montgomery Bertschy, Laura Healey, Zach Barrons, Wouter Hoogkamer; Experimental design: John Kuzmeski, Montgomery Bertschy, Zach Barrons, Wouter Hoogkamer; Data collection: John Kuzmeski, Montgomery Bertschy, Zach Barrons; Data analysis: John Kuzmeski, Montgomery Bertschy, Zach Barrons; Writing - original draft preparation: John Kuzmeski, Montgomery Bertschy, Laura Healey, Wouter Hoogkamer; Writing - review and editing: John Kuzmeski, Montgomery Bertschy, Laura Healey, Zach Barrons, Wouter Hoogkamer

